# Inhibitory synaptic transmission is impaired in the Kölliker-Fuse of male, but not female, Rett Syndrome Mice

**DOI:** 10.1101/2023.10.02.560501

**Authors:** Jessica R. Whitaker-Fornek, Paul M. Jenkins, Erica S. Levitt

**Author notes:** Corresponding Author: Jessica Whitaker-Fornek, 1150 W. Medical Center Dr., Ann Arbor, MI 48109-5632. Author Contributions: JW performed the experiments, analyzed data, prepared figures and wrote the manuscript. ESL conceptualized the study and assisted JW with experimental design, statistical analysis of data, revised and edited drafts of the manuscript and figures. PMJ provided expertise in axon initial segment imaging and revised manuscript and figures.

## Abstract

Rett Syndrome (RTT) is a severe neurodevelopmental disorder that mainly affects girls and women due to silencing mutations in the X-linked *MECP2* gene. One of the most troubling symptoms of RTT is breathing irregularity, including apneas, breath-holds, and hyperventilation. Mice with silencing mutations in *Mecp2* exhibit breathing abnormalities similar to human patients and serve as useful models for studying mechanisms underlying breathing problems in RTT. Previous work implicated the pontine, respiratory-controlling Kölliker-Fuse (KF) in the breathing problems in RTT. The goal of this study was to test the hypothesis that inhibitory synaptic transmission is deficient in KF neurons from symptomatic male and female RTT mice. We performed whole-cell voltage-clamp recordings from KF neurons in acute brain slices to examine pharmacologically isolated, spontaneous and electrically evoked inhibitory post-synaptic currents (IPSCs) in RTT mice and age- and sex-matched wild type mice. The frequency of spontaneous IPSCs was reduced in KF neurons from male RTT mice, but not female RTT mice. In addition, electrically evoked IPSCs were less reliable in KF neurons from male RTT mice, but not female RTT mice. KF neurons from male RTT mice were also more excitable and exhibited shorter duration action potentials. Increased excitability of KF neurons from male mice was not explained by changes in axon initial segment length. These findings indicate impaired inhibitory neurotransmission and increased excitability of KF neurons in male, but not female RTT mice, and suggest that sex-dependent mechanisms contribute to breathing problems in RTT.

**New and Noteworthy:** Kölliker-Fuse (KF) neurons in acute brain slices from male Rett syndrome (RTT) mice receive reduced inhibitory synaptic inputs compared with wild type littermates. In female RTT mice, inhibitory transmission was not different in KF neurons compared with controls. The results from this study show that sex-specific alterations in synaptic transmission occur in the KF of RTT mice.

**Graphical Abstract:** 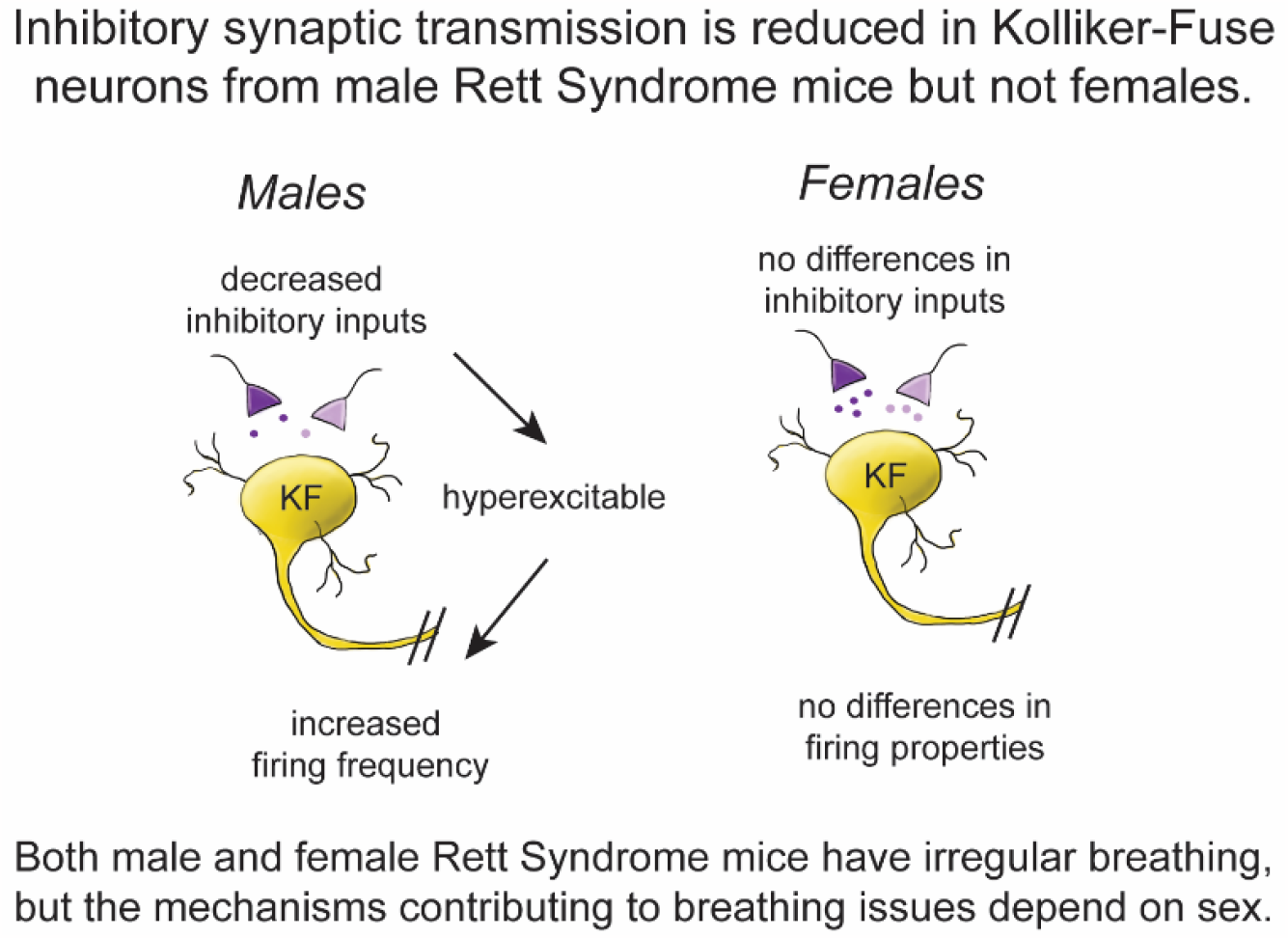

## Introduction

Rett Syndrome (RTT) is a rare neurodevelopmental disorder that mainly affects girls and women (Ivy & Standridge, 2021). One of the most concerning symptoms of RTT is unpredictable, irregular breathing. RTT patients exhibit breath holding, hyperventilation, apnea, apneusis and other breathing abnormalities (Ramirez et al., 2020; Ramirez et al., 2022). Breathing problems disrupt daily activities such as eating and physical therapy sessions and sometimes lead to sudden death (Kerr et al., 1997; Singh et al., 2020; Tarquinio et al., 2018; Weese-Mayer et al., 2006).

RTT is caused by silencing mutations in the X-linked gene *MECP2* that encodes methyl-CpG binding protein 2 (Amir et al., 1999). Like human RTT patients, mice with loss-of-function mutations in the *Mecp2* gene exhibit breathing abnormalities (Guy et al., 2001; Viemari et al., 2005). Respiratory disturbances are very severe in male RTT mice that completely lack Mecp2 protein (Katz et al., 2009; Voituron et al., 2009). Female RTT mice also have breathing problems that arise developmentally later compared to males, likely due to mosaic expression of functional Mecp2 protein as a result of partial X -inactivation (Abdala et al., 2010; Bissonnette & Knopp, 2008; Jiang et al., 2017; Levitt et al., 2013; Samaco et al., 2013). Male and female RTT mice are useful models for uncovering dysfunction within the brainstem respiratory control network that underlies breathing irregularity in RTT.

Respiratory-controlling Kölliker-Fuse (KF) neurons have been implicated in the breathing irregularities shown in male (Stettner et al., 2007) and female RTT mice (Abdala et al., 2010; Abdala et al., 2016). KF neurons are located in the dorsolateral pons where they contribute to the control of breathing and upper airway patency (Dutschmann & Dick, 2012; Dutschmann & Herbert, 2006; Varga et al., 2021). The KF supplies glutamatergic inputs to many downstream targets including respiratory rhythm and pattern generating neurons in the medulla (Geerling et al., 2017; Yokota et al., 2007). KF neuron activation enhances the post-inspiratory phase of breathing resulting in a transient apnea (Chamberlin & Saper, 1994; Paton & Dutschmann, 2002; Saunders & Levitt, 2020). These apneas are similar to what is observed in RTT mice (Abdala et al., 2010). Therefore, excessive excitatory or deficient inhibitory transmission onto KF neurons has been hypothesized to contribute to apneas in RTT mice. Indeed, enhancing GABA levels in the KF by microinjection of a GABA reuptake inhibitor decreased apneas observed in recordings of respiratory motor output from female RTT mice (Abdala et al., 2016). In addition, the number of GABAergic neurites is reduced in the KF of RTT mice (Abdala et al., 2016). Together, these results strengthen the hypothesis that respiratory-controlling KF neurons receive fewer inhibitory inputs in female RTT mice. While it has been shown that inhibitory synaptic transmission is impaired in other regions of the brainstem respiratory network from RTT mice (Chen et al., 2018; Medrihan et al., 2008; Xing et al., 2021), it remains unknown whether inhibitory synaptic transmission is deficient in the KF of male and female RTT mice.

The goal of this research was to examine inhibitory synaptic transmission and excitability in the KF of male and female RTT mice. We used acute brain slice electrophysiology to test the hypothesis that inhibitory synaptic transmission is reduced in the KF of male and female RTT mice. We found that inhibitory neurotransmission was impaired in male, but surprisingly not female RTT mice. These results suggest that multiple synaptic mechanisms may be involved in a sex-specific manner in the breathing abnormalities observed in RTT.

## Methods

### Animals

All experiments were approved by the University of Florida or the University of Michigan Institutional Animal Care and Use Committee (IACUC). All procedures using animals were in accordance with the National Institutes of Health Guide for the Care and Use of Laboratory Animals. Female heterozygous *Mecp2* knockout mutation mice, B6.129P2(C)-Mecp2^tm1.1Bird^ (stock no. 003890, Jackson Labs, Bar Harbor, ME) were purchased from Jackson Labs and backcrossed with male wild-type C57Bl/6J mice for three generations to establish the colony, then maintained by crossing wild-type males from the colony. 4-6 week old *Mecp2*^Bird/y^ mice (male RTT mice) and 6-10 month old *Mecp2*^Bird/+^ mice (female RTT mice) were used for all experiments. Hindlimb clasping was observed in all RTT mice used for the experiments in this study. Age and sex-matched wild type littermates were used as controls.

### Electrophysiology

Acute brain slices containing Kölliker-Fuse (KF) neurons were prepared as previously described (Levitt et al., 2015). Briefly, mice were anesthetized with isoflurane just prior to decapitation and brain removal. To acquire KF containing slices, the brain was blocked and mounted caudal end up in a vibratome chamber (Leica VT 1200S). Coronal slices (230 μm) were collected beginning at the caudal extent of the KF through the rostral extent of the KF. Brain slices were stored in warm (∼32 °C), oxygenated (95% O_2_, 5% CO_2_) artificial cerebrospinal fluid (aCSF) containing (in mM) 126 NaCl, 2.5 KCl, 1.2 MgCl_2_, 2.6 CaCl_2_, 1.2 NaH_2_PO_4_, 11 D-glucose and 21.4 NaHCO_3_. MK-801 (10 μM) was added to the cutting and slice storage solution to prevent excitotoxicity. Following at least 30 minutes of incubation, slices were placed into a recording chamber and perfused with warm (34°C) oxygenated aCSF with a flow rate of 1.5-3 ml/min.

Cells were visualized using an upright microscope (Nikon FN1) equipped with a custom-built IR-Dodt gradient contrast illumination and DAGE-MTI IR-2000 camera. KF neurons were located just ventral and lateral to the superior cerebellar peduncle.

Whole-cell voltage-clamp recordings were made from KF neurons using a Multiclamp 700B amplifier (V_hold_ = -60 mV). Glass recording pipettes (1.7 – 3 MΩ) were filled with high chloride internal solution containing (in mM): 115 KCl, 20 NaCl, 1.5 MgCl2, 5 HEPES(K), 2 BAPTA, 1-2 Mg-ATP, 0.2 Na-GTP, adjusted to pH 7.35 and 275-285 mOsM. The liquid junction potential (10 mV) was not corrected. Data were low-pass filtered at 10 kHz and collected at 20 kHz with pClamp 10.7 (Molecular Devices, Sunnyvale, CA), or collected at 400 Hz with PowerLab (LabChart version 5.4.2; AD Instruments, Colorado Springs, CO). Series resistance was monitored without compensation and remained <20 MΩ for inclusion. Current-clamp mode was used to record action potential firing following current injection (+200 pA, 500 ms step duration).

GABAergic and glycinergic IPSCs were isolated using the glutamate receptor blockers DNQX (10 μM) in the perfusate and pre-treatment with MK-801 (10 μM) in the cutting solution. Electrically evoked IPSCs (eIPSCs) were elicited using a bipolar stimulating electrode placed in the KF. Stimulation trials consisted of 20 sweeps of paired electrical pulses (0.2 ms duration, 50 ms interpulse interval) that were delivered every 20 seconds. Stimulation intensity (10-50 μA) was adjusted to evoke a submaximal postsynaptic current. In a subset of experiments, the glycine receptor antagonist strychnine (1 μM) and the GABA_A_ receptor antagonist gabazine (1 μM) were applied to ensure all currents were blocked.

We did not differentiate spontaneous release from action-potential independent release. In a subset of experiments, we added TTX (1 μM) and found that this had no effect on IPSC frequency. The average frequency of glycinergic sIPSCs (4 ± 0.82 Hz, n = 16) was not different than glycinergic miniature-post-synaptic currents (5 ± 1.1 Hz, n = 11) (unpaired t-test, p = 0.3849). The average frequency of GABAergic sIPSCs (8 ± 1.1 Hz, n = 18) was not different than GABAergic miniature-post-synaptic currents (8 ± 2 Hz, n = 10) (unpaired t-test, p = 0.7755). The majority of sIPSCs were likely action-potential independent.

### Drugs

All blockers were reconstituted and stored according to manufacturer’s instructions. (+)-MK-801 hydrogen maleate (Product No. M108), DNQX (Product No. D0540) and strychnine hydrochloride (Product No. S8753) were from Sigma-Aldrich. SR 95531 hydrobromide (Gabazine) was from Tocris Biosciences (Cat. No. 1262). Fresh solutions at the appropriate final concentrations were made fresh daily from concentrated stock solutions.

### Immunohistochemistry

Mice were deeply anesthetized with isoflurane and transcardially perfused with ice-cold phosphate-buffered saline (PBS) followed by 10% formalin. Brains were removed and placed in 10% formalin for 24 hours before immersion in 20% sucrose overnight and 30% sucrose for ∼6 h. Cryoprotected brains were frozen in optimal cutting temperature (O.C.T.) compound (Fisher Tissue Plus) on dry ice and stored at -80°C. Coronal brain slices (40 μm thick) containing the KF were cut using a Leica cryostat and stored as free-floating sections in PBS at 4°C. Brain slices were permeabilized using PBS containing 0.3% Triton-X-100 (PBS-T) and blocked for 1 hour with a buffer solution containing (3% normal goat serum and 2% bovine serum albumen) and incubated overnight at 4°C with primary antibody rabbit anti-ankyrin G (lab-generated, 1:1000) (Jenkins et al., 2013) diluted with blocking buffer. Brain slices were rinsed with PBS-T and incubated with secondary antibody goat anti-rabbit Alexa Fluor 647 (Invitrogen, AE32733, 1:500) diluted with blocking buffer for 1.5 hours at room temperature. Finally, slices were rinsed with PBS-T and mounted on slides with Fluoromount G with DAPI mounting media (Invitrogen). Brain slices containing KF neurons were imaged using a Zeiss LSM 880 confocal microscope equipped with Airyscan processing. Immunostaining experiments were performed in parallel on brain slices from six, 6-week-old male mice (n = 3 RTT, n = 3 WT littermates). The KF area from WT and RTT mouse brains (n = 6 mice; 3 images per mouse) were immunostained and imaged in parallel with a 63X NA1.4 Oil/DIC Plan-Apochromat objective and 633 nm lasers. Images were pseudocolored white.

### Axon initial segment quantification

Z stacks (20 μm, 10 slices) were collapsed to create maximum intensity z projections in Fiji (Schindelin et al., 2012). The segmented line measure tool was used to acquire a fluorescence profile along the length of the axon initial segment (AIS). The start and end of the AIS was defined as 15% of the maximum fluorescence value. AIS length was measured as the distance between the start and end of the AIS. Normalized (*i*.*e*., background subtracted) mean fluorescence intensity for the total ankyrin-G signal was calculated in Fiji for each maximum intensity z projection taken of the KF area (n = 8 images from 3 RTT mice, n = 9 images from 3 wild type littermates).

### Data analysis

Spontaneous IPSCs were detected and analyzed for 5-minute epochs during baseline and following drug exposure (Clampfit 11.1, Molecular Devices). For electrically-evoked IPSCs, the average amplitude and decay tau of the first eIPSC of the pair was calculated from the averaged waveform for 20 sweeps. Paired-pulse ratio was determined by dividing the average amplitude of the second eIPSC by the average amplitude of the first eIPSC (P2/P1). The failure rate for the first eIPSC of the pair (*i*.*e*., the percentage of stimulations that failed to evoke a postsynaptic current) was calculated by dividing the number of failed stimulations by the total number of stimuli (20) and multiplying by 100. Average action potential firing frequency was calculated using the action potential search mode in Clampfit. The average action potential waveform per neuron was used to calculate duration from peak to after-hyperpolarization, rise time (75 to 15% of peak) and decay time (10% to 90% of peak) in Clampfit.

All statistical analyses were performed using Graphpad Prism (version 9). All error bars represent SEM unless otherwise stated. Replicates are individual neurons. 1-2 neurons were recorded per animal. Comparisons between two groups were made using unpaired t-test or Mann-Whitney U test depending on the results of normality testing (D’Agostino-Pearson).

## Results

To examine inhibitory synaptic transmission in the respiratory-controlling KF area of symptomatic male and female RTT mice, we made whole-cell voltage-clamp recordings from KF neurons contained in acute brain slices from male hemizygous *Mecp2*^*Bird/y*^ mice (male RTT mice), female heterozygous *Mecp2*^*Bird/+*^ mice (female RTT mice) and their age- and sex-matched wild type littermates. Male hemizygous *Mecp2*^Bird/y^ mice (male RTT mice) are null for Mecp2 function, given *Mecp2* is X-linked. Males exhibit severe respiratory and motor deficits by 4 weeks of age, with a lifespan of 50-60 days (Viemari et al., 2005). Therefore, 4-6 week-old male RTT mice were used for all experiments. Female, heterozygous *Mecp2*^Bird/+^ mice (female RTT mice) display respiratory problems at 6 months of age or older (Jiang et al., 2017).

Therefore, 6-10 month-old adult female mice were used for all experiments.

### Spontaneous inhibitory post-synaptic currents are less frequent in KF neurons from male RTT mice

We measured spontaneous inhibitory post-synaptic currents (sIPSCs) from KF neurons in brain slices from male RTT mice and age-matched wild type littermate controls (Fig. 1, A). The frequency of sIPSCs was significantly reduced in male RTT mice compared with wild-type mice (Fig. 1, B; p = 0.0012, unpaired t-test). However, the amplitude of sIPSCs was not different for KF neurons from male RTT mice compared to wild type littermates (Fig. 1, C; p = 0.1151, unpaired t-test). Differences in the decay kinetics of sIPSCs often indicates differences in receptor subunit composition (Bosman et al., 2002). There was no difference in the decay tau for sIPSCs from KF neurons from male RTT mice compared to WT controls (Fig. 1, D; p = 0.8631, Mann-Whitney Test). In a subset of experiments, co-application of gabazine (1 μM) and strychnine (1 μM) eliminated all sIPSCs (average frequency = 0 Hz, n = 4 cells from 3 animals), confirming IPSCs were due to GABA_A_ and glycine receptor activation, respectively.

**Figure 1.**
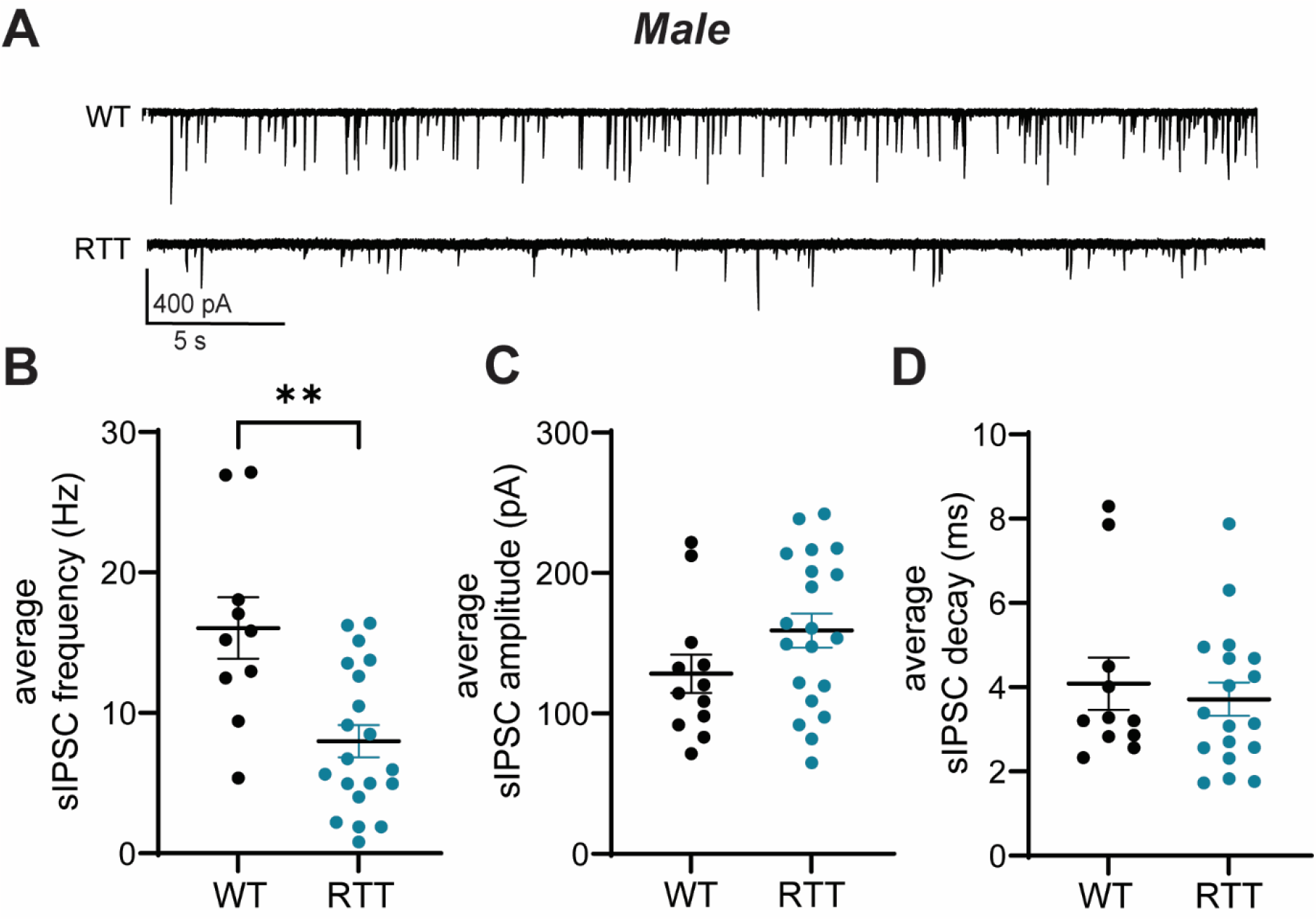
Spontaneous inhibitory synaptic transmission in KF neurons from male RTT mice. A: Representative traces of spontaneous inhibitory postsynaptic currents (sIPSCs) recorded from KF neurons contained in brain slices from male Rett Syndrome (RTT) mice and age-matched wild type (WT) littermates. B: The frequency of sIPSCs was reduced in KF neurons from RTT mice compared to controls (**, p = 0.0012, unpaired t-test). C: There was no difference in the amplitude of sIPSCs (p = 0.1151, unpaired t-test). D: The average decay tau of sIPSCs was not different for KF neurons from male RTT mice compared to controls (p = 0.8631, Mann-Whitney test). Line and error bars indicate mean ± SEM. Individual data points are from individual neurons, 1-2 neurons per mouse.

### Spontaneous inhibitory post-synaptic currents are not different in KF neurons from female RTT mice

To examine inhibitory synaptic transmission in the KF of female RTT mice, we performed whole-cell voltage clamp recordings from KF neurons contained in brain slices from female RTT mice and age-matched wild type littermate controls (Fig. 2, A). In KF neurons from WT female mice, the average frequency of sIPSCs was lower compared to WT males (p = 0.0023, unpaired t-test). In contrast to male RTT mice, the frequency of sIPSCs was not different for RTT females compared to wild type littermates (Fig. 2, B; p = 0.4668, Mann-Whitney test). The amplitude of sIPSCs was also not different for KF neurons from female RTT mice compared to wild type controls (Fig. 2, C; p = 0.4018, unpaired t-test). The decay tau was also not different for sIPSCs female RTT mice compared with wild type littermates (Fig. 2, D; p = 0.2918, Mann-Whitney Test). These results suggest that spontaneous inhibitory transmission is not impaired in KF neurons from female RTT mice.

**Figure 2.**
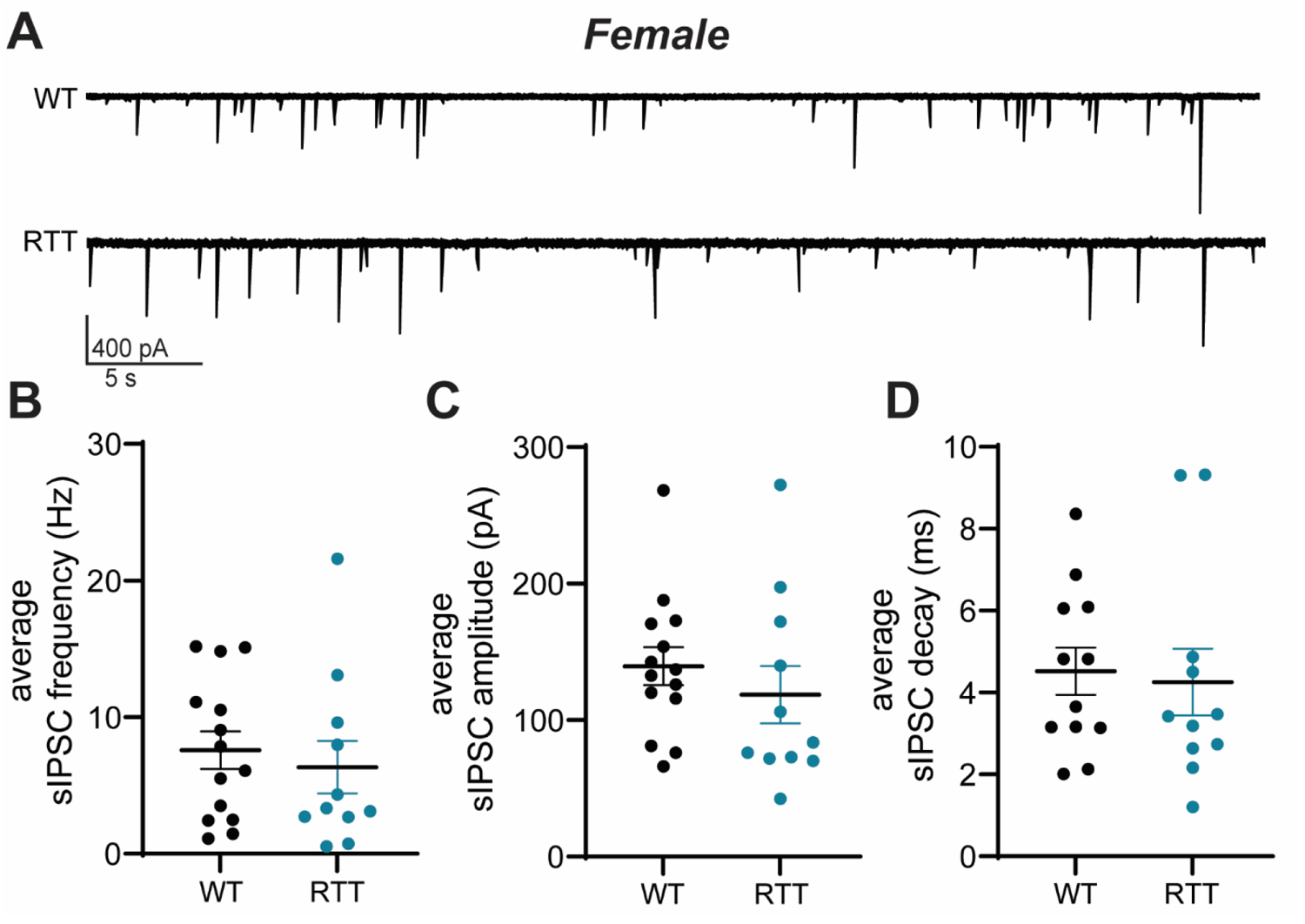
Spontaneous inhibitory synaptic transmission in KF neurons from female RTT mice. A: Representative traces of spontaneous inhibitory postsynaptic currents (sIPSCs) recorded from KF neurons contained in brain slices from female RTT mice and age-matched WT littermates. B. In contrast to males, sIPSC frequency was not different in female RTT mice compared to controls (p = 0.4668, Mann-Whitney test). C. The average sIPSC amplitude was not different in female RTT mice compared to controls (p = 0.4018, unpaired t-test). D. The average sIPSC decay tau was not different for female RTT mice compared to controls (p = 0.2918, unpaired t-test). Line and error bars indicate mean ± SEM. Individual data points are from individual neurons, 1-2 neurons per mouse.

### Evoked inhibitory synaptic currents in KF neurons from male mice

To investigate responses to stimulated GABA and glycine release in the KF, we delivered paired electrical stimuli to the KF and measured evoked inhibitory postsynaptic currents (eIPSCs) in KF neurons from male RTT mice and wild type littermates (Fig. 3, A). There was no difference in eIPSC amplitude for KF neurons from male RTT mice compared to wild type controls (Fig. 3, B; p = 0.1453, unpaired t-test). There was also no difference in the decay tau of eIPSCs for KF neurons from male mice (Fig. 3, C; p = 0.3482, unpaired t-test). In a subset of experiments, we confirmed that co-application of strychnine (1 μM) and gabazine (1 μM) blocked electrically evoked currents (average peak amplitude: 10.8 ± 2.9 pA, n = 4 cells, from 3 animals).

**Figure 3.**
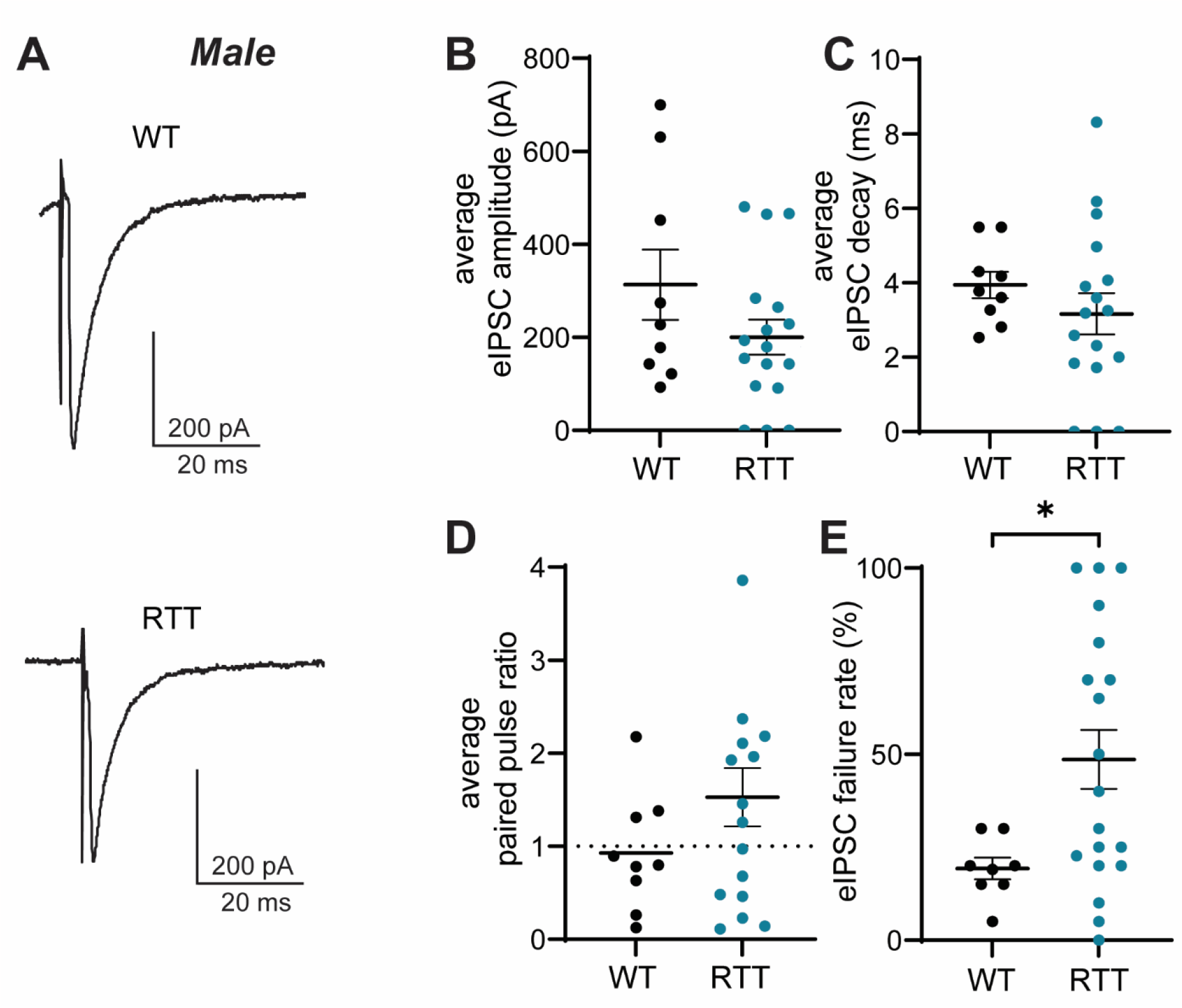
Evoked inhibitory synaptic transmission in KF neurons from male RTT mice. A: Representative traces of electrically evoked inhibitory postsynaptic currents (eIPSCs) recorded from KF neurons from male RTT mice and WT controls. The average eIPSC amplitude (B), decay tau (C) and paired pulse ratio (D) for male RTT mice were not different from controls (amplitude: p = 0.1453; decay tau: p = 0.3482, paired-pulse ratio: p = 0. 1957, unpaired t-tests). E: The percentage of stimulations that failed to evoke an IPSC was higher in male RTT mice compared to controls (*, p = 0.0219, unpaired t-test). Line and error bars indicate mean ± SEM. Individual data points are from individual neurons, 1-2 neurons per mouse.

To examine differences in the release properties of GABA and glycine onto KF neurons from wild type and RTT male mice, we calculated the paired pulse ratio (average amplitude of the eIPSC for pulse 2/average amplitude of the eIPSC for pulse 1) (Fig. 3, D). Both paired pulse facilitation and paired pulse depression were observed for eIPSCs. There was no difference in paired pulse ratio for eIPSCs from male RTT mice compared to controls (p = 0.1957, unpaired t-test).

Electrical stimulation did not always evoke an IPSC on every sweep. We determined the percentage of stimulations that failed to evoke an IPSC by dividing the number of failed stimulations by the total number of sweeps (Fig. 3, E). In wild type mice, approximately 20% of stimulations failed to evoke an IPSC (Fig. 3, E). There were significantly more failures for KF neurons from male RTT mice compared to controls (p = 0.0219, unpaired t-test), despite the fact that stimulation intensity did not differ between genotypes (p = 0.3702, Mann-Whitney test). The increased eIPSC failure rate in male RTT mice is consistent with impaired presynaptic release properties.

### Evoked inhibitory synaptic currents in KF neurons from female mice

Electrically evoked IPSCs (eIPSCs) were recorded from KF neurons in brain slices from female RTT mice and age-matched wild type littermates (Fig. 4, A). There was no difference in the amplitude of eIPSCs in KF neurons from female RTT mice compared to wild type controls (Fig. 4, B; p = 0.7939, unpaired t-test). The decay tau of eIPSCs was also not different for female RTT mice compared to wild type controls (Fig. 4, B; p = 0.2595, unpaired t-test). Similar to males, there was no difference in stimulus intensity used to evoke IPSCs in KF neurons from female RTT mice compared to controls (p = 0.4105, Mann-Whitney test).

**Figure 4.**
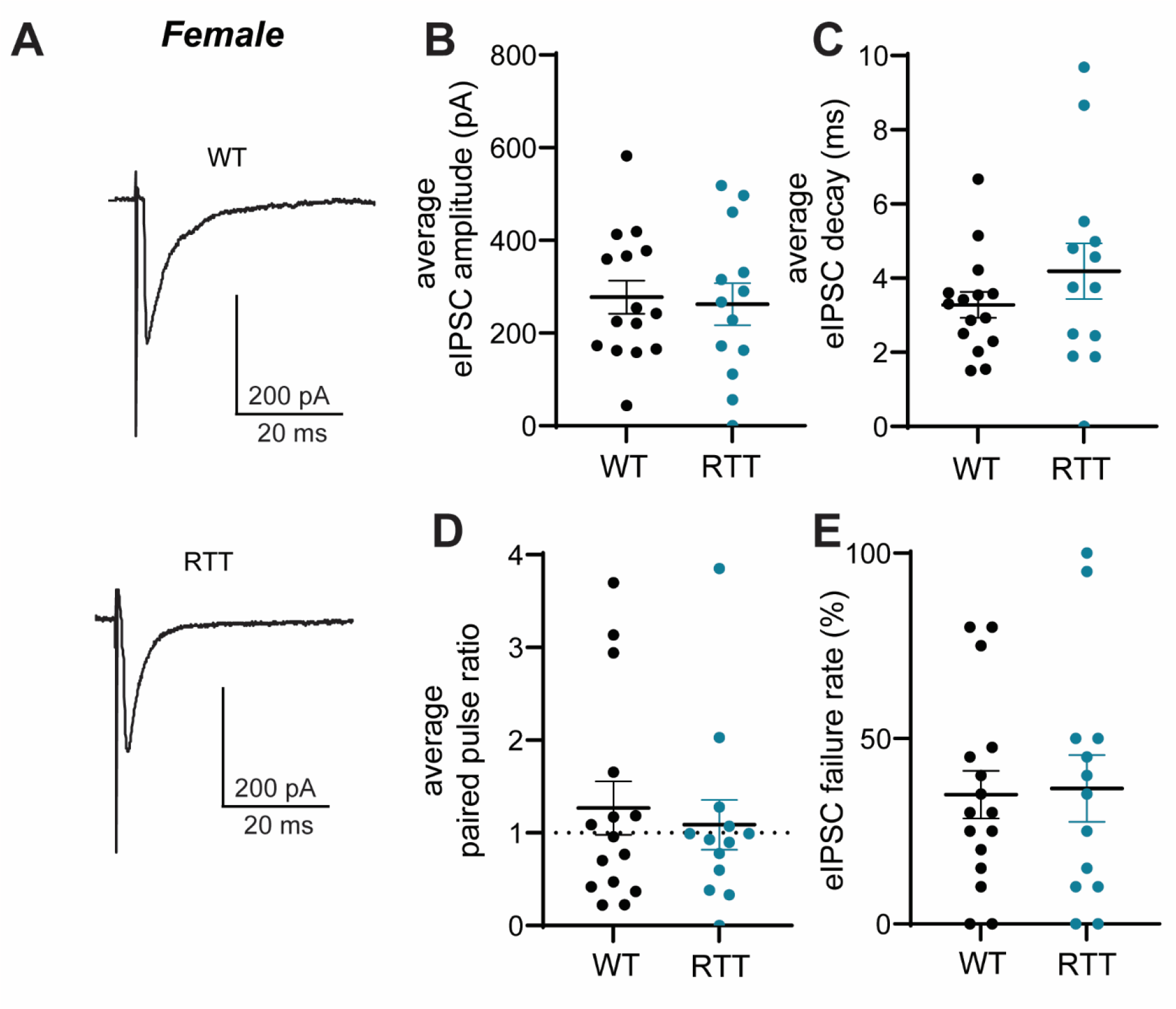
Evoked inhibitory synaptic transmission in KF neurons from female RTT mice. A: Representative traces of electrically evoked inhibitory postsynaptic currents (eIPSCs) recorded from KF neurons from female RTT mice and WT controls. Summarized data for the average eIPSC amplitude (B), decay tau (C) for female RTT mice were not different from controls (amplitude: p = 0.7939; decay tau: p = 0.2595; unpaired t-tests). D: The average paired pulse ratio was not different for female RTT mice compared to controls (p = 0.8562, Mann-Whitney test). E. In contrast to males, no significant differences were observed for the percentage of failed eIPSCs in female RTT mice compared to WT controls (p = 0.8966, unpaired t-test). Line and error bars indicate mean ± SEM. Individual data points are from individual neurons, 1-2 neurons per mouse.

Paired pulse ratio and the percentage of failures were examined as an indicator of GABA and glycine release properties. Similar to males, eIPSCs showed both paired-pulse facilitation and depression (Fig. 4, D) and there was no difference in paired-pulse ratio (p = 0.8562, Mann-Whitney Test). Failures to evoke IPSCs were also observed in KF neuron recordings from female RTT and wild-type mice (Fig. 4, E). However, there was no difference in the percentage of failures for eIPSCs in KF neurons from female mice (p = 0.8966, unpaired t-test). Thus, unlike males, there were no differences observed in electrically evoked release properties of inhibitory neurotransmitter in the KF of female RTT mice compared to wild-type controls.

### Action potential properties of KF neurons from male mice

The excitability of KF neurons in response to depolarizing current injections were examined at the beginning of all experiments. KF neurons from both male RTT mice and wild type littermates fired action potentials as a result of depolarizing current injections (200 pA, 500 ms) (Fig. 5, A). KF neurons from male RTT mice fired action potentials at a higher average frequency compared to WT littermates (Fig. 5, B, p = 0.0413, Mann-Whitney Test). Consistent with higher frequency firing, the duration of action potentials was shorter for KF neurons from male RTT mice compared to controls (Fig. 5, C, p = 0.0096, Mann-Whitney Test). Although the rise time for action potentials was not different (Fig. 5, D, p = 0.0726, Mann-Whitney Test), the difference in duration was due to faster decay times in KF neurons from male RTT mice (Fig.5, E, p = 0.0039, Mann-Whitney test).

**Figure 5.**
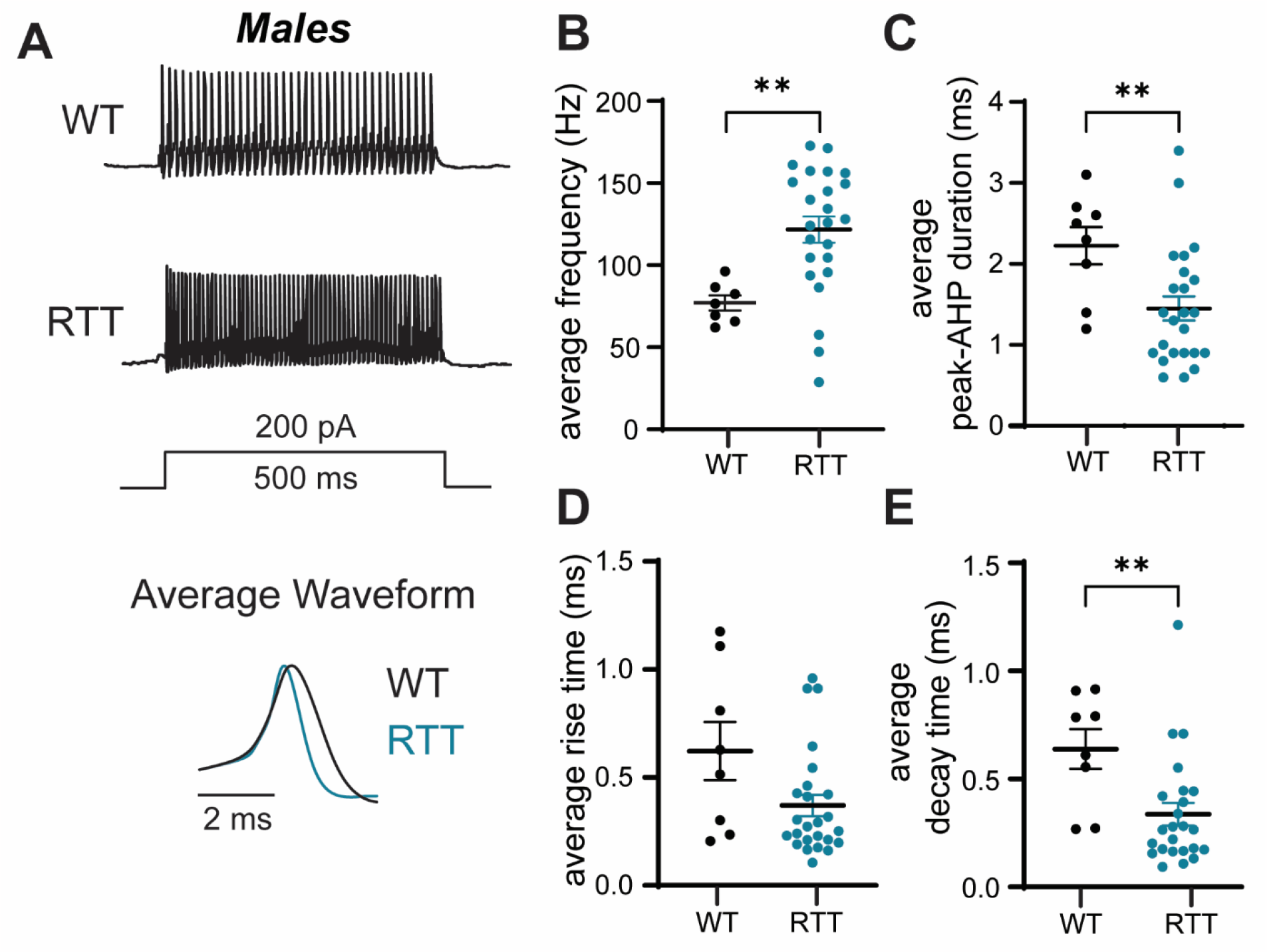
Firing properties of KF neurons from male RTT mice. A: Representative whole cell current clamp recordings from KF neurons from a WT male mouse (top trace) and a RTT male mouse (bottom trace). Average action potential waveforms are overlaid below. B-E: Summary of average action potential firing frequency (B), action potential duration measured from the peak to after-hyperpolarization trough (C), average rise time (D) and average decay time (E). KF neurons from male RTT mice fired higher frequency action potentials compared to wild type controls (B) (p = 0.0413, Mann-Whitney test). Average action potential duration was shorter for KF neurons from RTT mice (C) (p = 0.0096, Mann-Whitney test). Action potentials fired by KF neurons from RTT mice did not have significantly faster rise times (D) (p = 0.0726, Mann-Whitney test) but decay times were significantly faster compared to controls (E) (p = 0.0039, Mann-Whitney test). Line and error bars indicate mean ± SEM. Individual data points are from individual neurons, 1-2 neurons per mouse.

### Axon initial segment length in KF area from male mice

Neuronal excitability and firing properties are influenced by structural properties of the axon initial segment (AIS) (Yamada & Kuba, 2016). Therefore, we immunofluorescently labeled ankyrin-G, the scaffolding protein responsible for organization of the ion channels necessary for action potential initiation at the AIS (Nelson & Jenkins, 2017), to examine the AIS length in KF neurons from male RTT mice and wild type littermates (Fig. 6, A). We found there was no difference in average AIS length for KF neurons from male RTT mice compared to controls (Figure 6, B, p = 0.2398, Mann-Whitney test). There was also no effect of genotype on the mean fluorescence intensity of ankyrin-G immunostaining in the KF area (p = 0.5159, unpaired t-test, n = 6 mice (3 WT, 3 RTT), 2-3 images per mouse).

**Figure 6.**
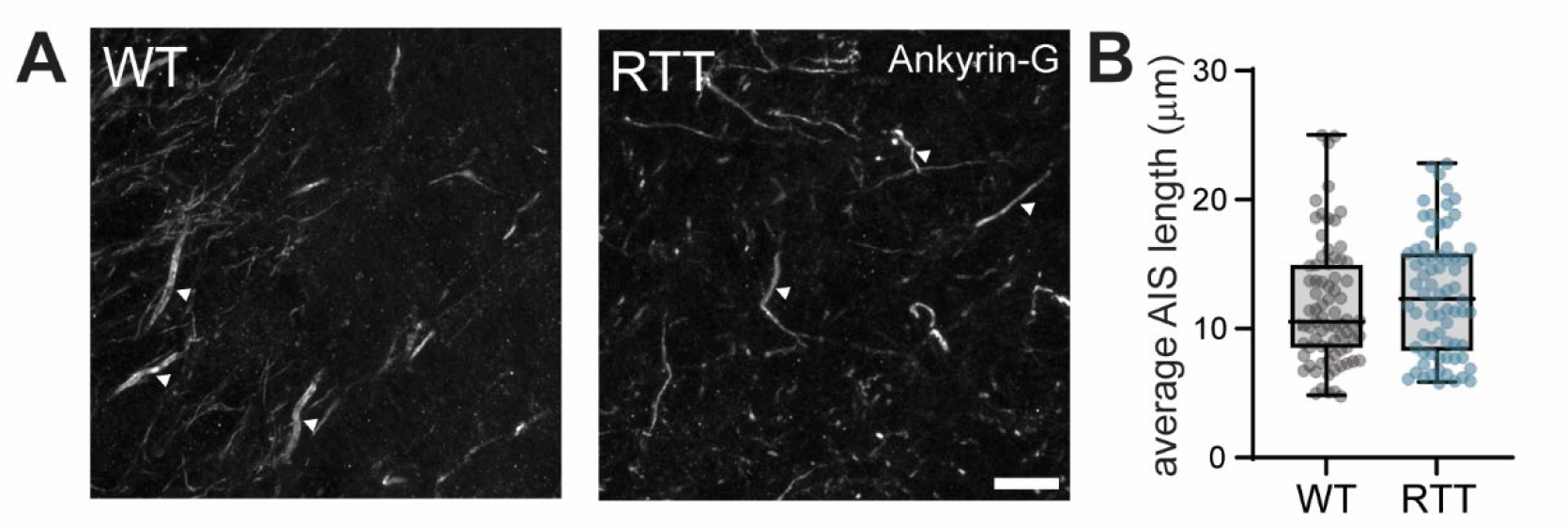
Axon initial segments in the KF area from male RTT mice. A. Representative maximum intensity z projection images (60x magnification) showing ankyrin-G immunostaining in the KF area from a male WT and male RTT mouse. Ankyrin-G staining was used to visualize axon initial segments (AIS; white arrows denote three examples per image). Scale bar = 10 μm. B. Average length of the AIS was not different for male RTT mice compared to WT controls (p = 0.2398, Mann-Whitney test, n = 6 mice (3 WT, 3 RTT), 2-3 images per mouse). Box and whisker plots indicate median and quartile ranges including the minimum and maximum values for individual segments (individual data points).

### Action potential properties of KF neurons from female mice

Similar to males, action potentials were elicited in KF neurons from female RTT mice and wild type littermates in response to depolarizing current (500 ms duration, 200 pA) (Fig. 7, A). In contrast to males, there were no differences in action potential firing frequency for KF neurons from female RTT mice compared to wild type littermates (Fig. 7, B, p = 0.7373, unpaired t test). There were also no differences in action potential duration (Fig. 7, C, p = 0.6492, Mann Whitney test), average rise time (Fig. 7, D, p = 0.3929, Mann-Whitney test), or average decay time (Fig. 7, E, p = 0.9680, Mann-Whitney test) for KF neurons from female RTT mice compared to controls.

**Figure 7.**
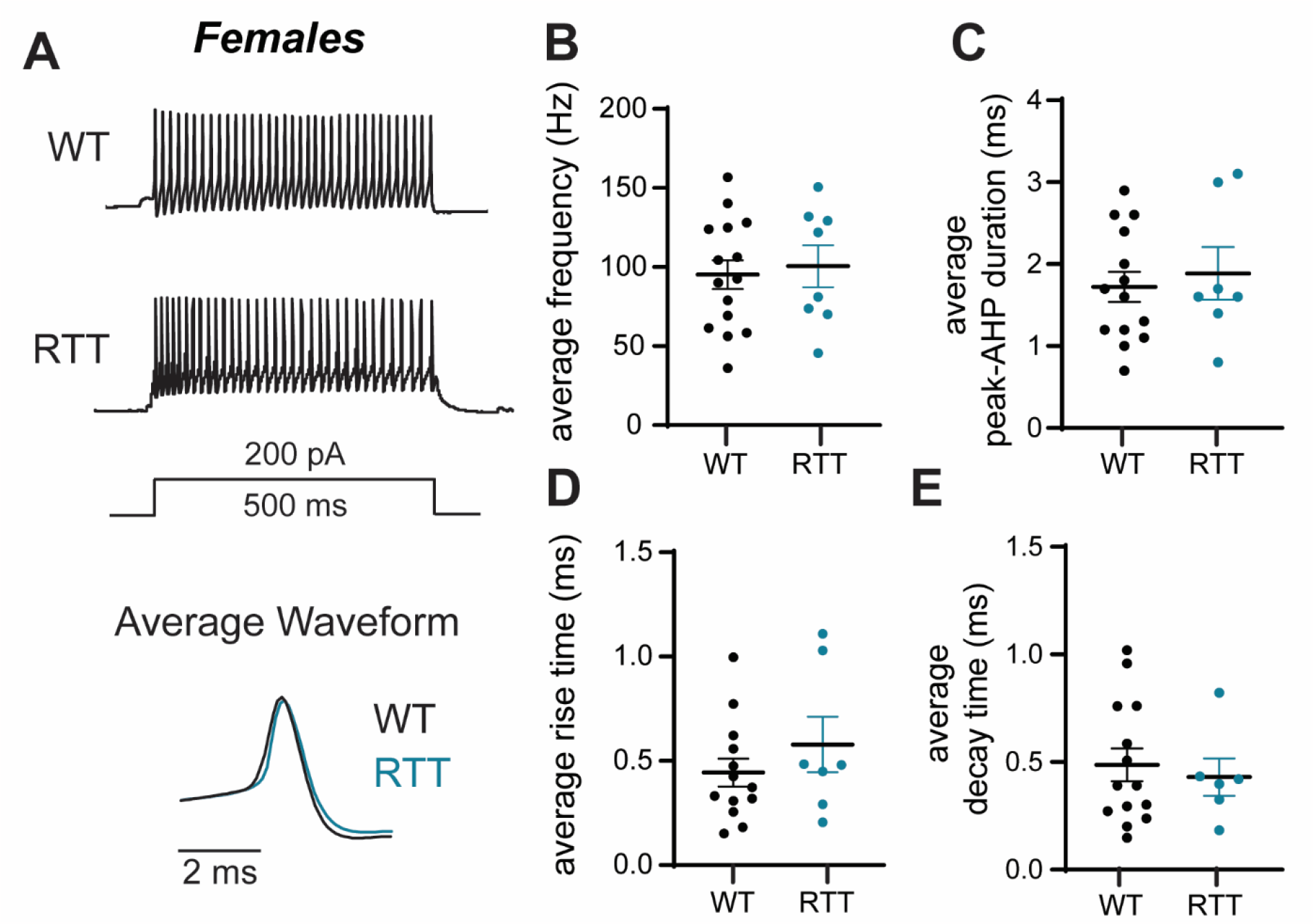
Firing properties of KF neurons from female RTT mice. A: Representative whole cell current clamp recordings from KF neurons from a WT female mouse (top trace) and a RTT female mouse (bottom trace). The average action potential waveforms are overlaid below. B-E: Summary of average action potential firing frequency (B), action potential duration measured from the peak to after-hyperpolarization trough (C), average rise time (D) and average decay time (E). Action potentials fired by KF neurons from female RTT mice were no different than wild type controls. There was no difference in the frequency of action potentials compared to wild type controls (B) (p = 0.7373, unpaired t test). Average action potential duration was not different for KF neurons from RTT mice (C) (p = 0.6492, Mann Whitney test). Also, there were no differences in the rise time (p = 0.3929, Mann Whitney test) or decay time (p = 0.9680, Mann-Whitney test) for action potentials fired by KF neurons from RTT mice. Line and error bars indicate mean ± SEM. Individual data points are from individual neurons, 1-2 neurons per mouse.

## Discussion

This study examined spontaneous and evoked inhibitory synaptic transmission in KF neurons from symptomatic male and female RTT mice in comparison to their wild type, age- and sex-matched littermates. In male RTT mice, we show that KF neurons exhibit deficient inhibitory synaptic transmission and are hyperexcitable. Deficient inhibitory synaptic transmission appears to be due to deficits in neurotransmitter release or synaptic number, since the frequency, but not amplitude, of spontaneous IPSCs was reduced and stimulation evoked IPSCs were less reliably elicited. However, in female RTT mice, we show there are no differences in the synaptic inhibitory transmission onto KF neurons or action potential firing properties of KF neurons compared to control age-, and sex-matched wild type littermates. This is an important sex difference since RTT primarily affects females, but much of the work on synaptic transmission in RTT mouse models have used male mice (Chen et al., 2018; Li, 2022; Medrihan et al., 2008).

Our findings for KF neurons from male RTT mice add to growing evidence in support of impaired inhibitory synaptic transmission within the brainstem breathing control network of male RTT mice (Li, 2022). GABAergic/glycinergic sIPSC frequency is also decreased in rostroventrolateral medullary neurons and nucleus of the solitary tract neurons from male RTT mice (Chen et al., 2018; Medrihan et al., 2008). We also show that KF neurons from male RTT mice fire higher frequency action potentials which is similar to the hyperexcitability present in medullary respiratory neurons (Wu et al., 2019) and mesencephalic trigeminal neurons (Oginsky et al., 2017) from male RTT rodent models. Hyperexcitability can occur due to structural plasticity of the axon initial segment where action potentials are initiated (Yamada & Kuba, 2016). Increased length of axon initial segments is found in hippocampal neurons in a mouse model of Angelman Syndrome (Kaphzan et al., 2011; Nelson & Jenkins, 2017). However, differences in axon initial segment length did not explain the hyperexcitability of KF neurons from male RTT mice in our study. Other properties of the AIS, such as increased voltage-gated potassium channel expression (Shu et al., 2007), may contribute to the shorter duration of action potentials we observed in KF neurons from male RTT mice.

Our results from female mice are unexpected given previously published data from Abdala and colleagues, who showed that the number of GABAergic neurites are reduced in the KF area of female RTT mice compared to wild type littermates (Abdala et al., 2016). In addition, blocking GABA reuptake locally within the KF decreases apneas in the in situ working heart brainstem preparation of female RTT mice (Abdala et al., 2016). Yet, we found no physiological evidence in support of deficient GABAergic synaptic transmission in the KF of female RTT mice. The different findings can be explained by the approaches we used to study inhibitory synaptic transmission onto KF neurons. First, we measured inhibitory synaptic transmission onto KF neurons directly using ex vivo electrophysiology, rather than indirectly by visualizing GABAergic projections using green fluorescent protein expression driven by the Gad67 promoter which is an anatomical proxy for GABA producing neurons. Second, an alternate explanation for the positive effects of the GABA reuptake inhibitor (Abdala et al., 2016) could be restoration of excitatory/inhibitory balance that was disrupted due to excessive excitatory transmission, rather than deficient inhibitory transmission, in the KF of RTT mice. To test this hypothesis, future experiments could measure spontaneous excitatory postsynaptic currents in KF neurons from female RTT mice.

### Limitations

The goal of the present study was to examine inhibitory synaptic currents in symptomatic male and, importantly, female RTT mice. Male RTT mice show breathing problems as early as 4-6 weeks old while female RTT mice show breathing irregularities once they reach 6 months old (Jiang et al., 2017). The substantial age difference between male and female symptomatic mice complicates direct comparisons between the sexes in the present work. For example, we cannot rule out age as a contributing factor to lower sIPSC frequency in KF neurons from WT female mice compared to WT males (Rozycka & Liguz-Lecznar, 2017). Future work examining synaptic transmission in female mice at pre-symptomatic ages could be informative to determine whether deficits are present earlier in development.

Second, we did not identify whether KF neurons from female RTT mice contained functional Mecp2 protein. Due to partial X-inactivation, approximately half of KF neurons in female RTT mice are Mecp2-negative and half are Mecp2-positive (Abdala et al., 2016). Both Mecp2-positive and Mecp2-negative KF neurons from female RTT mice were surrounded by significantly fewer GABAergic perisomatic bouton-like puncta compared to wild type littermates (Abdala et al., 2016). Therefore, while cell-autonomous effects are a possibility for some post-synaptic responses in RTT mice (Gantz et al., 2011), we do not think that cell-autonomous effects of Mecp2 expression in the post-synaptic neuron influenced inhibitory synaptic transmission onto KF neurons from female RTT mice in our study.

## Conclusion

Breathing problems can be life-threatening for girls and women diagnosed with RTT. Uncovering the neural dysfunction in the brainstem respiratory control network of RTT mice is a critical first step toward learning how to correct breathing problems in RTT patients. Using acute brain slice electrophysiology, we found that inhibitory synaptic transmission was impaired in the KF area of male RTT mice. Meanwhile, female RTT mice did not have decreased inhibitory transmission upon their KF neurons. The widespread imbalance of excitatory and inhibitory signaling in the brains of RTT mice poses a daunting challenge for restoring normal brain function and correcting the devastating neurological issues in human RTT patients. Targeted approaches are needed to restore excitation/inhibition balance across the breathing control network with the hope of improving the breathing abnormalities in RTT.

## Acknowledgements

This work was funded by the International Rett Syndrome Foundation Basic Research Grant #3608 to ESL. Thank you to Nicole Filipelli for technical assistance.

